# Differential HIV-1 Proviral Defects in Children vs. Adults on Antiretroviral Therapy

**DOI:** 10.1101/2025.05.23.655786

**Authors:** Jenna M. Hasson, Mary Grace Katusiime, Adam A. Capoferri, Michael J. Bale, Brian T. Luke, Wei Shao, Mark F. Cotton, Gert van Zyl, Sean C. Patro, Mary F. Kearney

## Abstract

HIV-1 proviral landscapes were investigated using near full-length HIV single-genome sequencing on blood samples from 5 children with vertically acquired infection and on ART for ∼7-9 years. Proviral structures were compared to published datasets in children prior to ART, children on short-term ART, and adults on ART. We found a strong selection for large internal proviral deletions in children, especially deletions of the *env* gene. Only 2.5% of the proviruses were sequence-intact, lower than in the comparative datasets from adults. Of the proviruses that retained the *env* gene, >80% contained two or more defects, most commonly stop codons and/or *gag* start mutations. Significantly fewer defects in the major splice donor site (MSD) and packaging signal were found in the children on short or long-term ART compared to the adults, and *tat* was more frequently defective in children. These results suggest that different selection pressures shape the proviral landscape in children compared to adults and reveal potentially different genetic regions to target for measuring the intact HIV reservoir and for achieving HIV remission in children.

## INTRODUCTION

Human immunodeficiency virus type-1 (HIV-1) in infants and children remains a significant global health challenge despite effective antiretroviral therapy (ART) for prevention and treatment. In 2023, 16% of pregnant women living with HIV-1 had no access to ART [1, 2], and this number could be rising today. Currently, approximately 1.5 million children aged 0-14 years are living with HIV-1, of which, only 57% had access to ART in infancy [1]. Without ART, 50% of children born with HIV-1 typically die by 2 years of age [3].

As there is currently no scalable cure, life-long ART adherence is required for children with HIV-1. In adults living with HIV-1, non-AIDS-related morbidities and drug toxicities have been observed in those on ART [4-6]. However, little is known about the long-term effects of HIV-1 and ART in children. HIV-1 reservoirs (the intact proviruses that persist on ART) in children with perinatally acquired infection are influenced by maternal ART and immune status during pregnancy, early life immune dynamics, and timing of ART initiation [7-10]. Current treatment guidelines recommend that infants perinatally exposed to HIV-1 should begin combination ART within 6 hours of birth [11, 12]. Very early ART initiation limits the size of the viral reservoir, as reservoir seeding occurs during uncontrolled viral replication [13-17]. Infants who initiate ART before 6 weeks of age have smaller proviral reservoirs than those treated later [17-21]. Proviral DNA levels decay following ART initiation until reaching a stable set point [19, 22-29], although a new case study suggests that reservoir levels may decay further after decades of ART in some individuals born with the virus [20].

Previous studies have investigated HIV-1 dynamics in a subset of children participating in the Children with HIV Early antiRetroviral (CHER; NCT00102960) trial [30]. ART was initiated at a median age of 5 months and viremia was suppressed for a median of 8 years at sampling. Van Zyl *et al.* found no evidence of viral replication while on ART in the children [31] and Katusiime *et al.* reported that only about 1% of persisting proviruses on long-term ART were genetically intact [32], a lower proportion than in most adults on long-term ART [33-35]. Integration site analyses demonstrated that infected cell clones were present in this cohort as early as 1.8 months of age and persisted for at least 9 years on ART [36]. It was also demonstrated that the largest infected cell clones were stable over time and contained solo HIV LTRs with no internal HIV sequences [37]. Additional pediatric studies have described proviral structures in pre-ART and during short-term ART and found that intact proviruses declined from ∼50% to ∼20% over the early months of treatment [38]. In addition, ART initiation prior to one week of age was associated with lower reservoir sizes [26].

Our current study addresses an important gap in the field, characterizing the structures of proviruses with internal HIV sequences that persist in children who adhere to long-term ART. Although previous studies have shown that the largest infected cell clones have no internal HIV sequences (*i.e.*, only solo LTRs), comprising about 50-90% of the infected PBMC in children on long-term ART, and that only about 1% of the infected cells contain inferred sequence-intact proviruses, the structures of proviruses in the remaining infected cells remains unknown. Characterizing the proviruses that contain HIV-1 gene-coding sequences on long-term ART may inform of the potential selection pressures that eliminate some infected cells while allowing others to persist, ultimately revealing new targets for eliminating the HIV-1 reservoir in children. In-depth characterization of the HIV-1 reservoir, as well as the defective proviral population that persists on ART, is vital to identifying the sources of the viral reservoir and to inform future curative strategies in children. Here, we investigated the genetic structures of proviruses persisting in children on long-term ART and compare them to proviral structures present in children prior to ART initiation, on short-term ART, and to those that persist in adults [33-35, 38].

## RESULTS

### Fewer proviruses in children on long-term ART are sequence intact, compared to children on short-term ART

A total of 882 (range 56 - 450) near full-length (NFL) proviral sequences were obtained from the 5 children in the CHER cohort who had viremia sustainably suppressed on ART for median 8 years (**Table 2**) (Genbank Accession: PV625462-PV626342). Of these sequences, 860 were defective and 22 (2.5%) were inferred intact using the ProSeq-IT multi-parameter intactness tool [39]. We compared the 882 proviral structures in children on long-term ART (7.4-8.8 years) to previously published NFL proviral sequence datasets obtained from children with varying times of ART initiation and durations of treatment (**Table 3**). These comparative datasets included proviruses in infants prior to ART initiation (n=132 sequences) (EIT study, pre-ART) [38], in children on short-term ART (<1 year) after initiating treatment <5 days after birth (n=244 sequences) (EIT study, on-ART) [38], and in children on short-term ART (<2.5 years) after receiving delayed treatment (<1 year old) (n=132 sequences) (EIT control group, delayed ART) [38]. We also compared the proviral structures from children on long-term ART to proviral structures in adults on ART for a median of 9.26 years (n=1,056 sequences) [33-35].

**Table 1.**
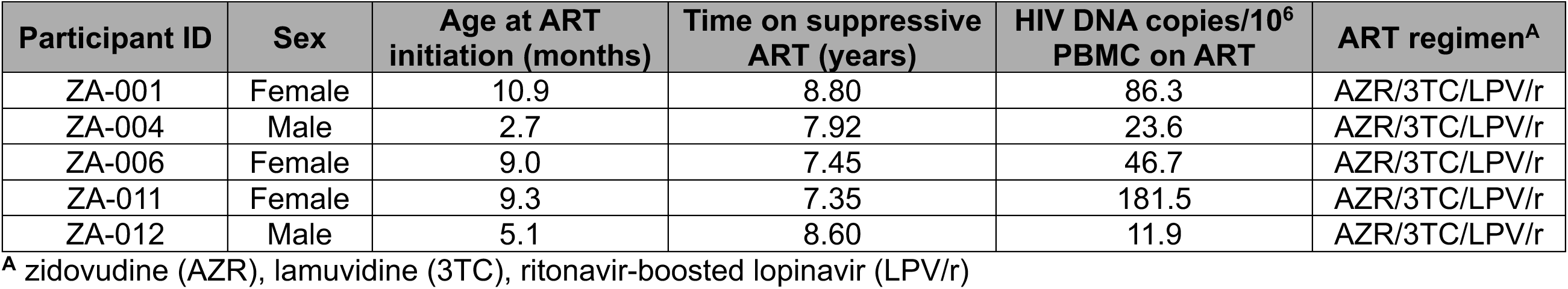
Participant demographic and virologic characteristics.

| **Participant ID** | **Sex** | **Age at ART initiation (months)** | **Time on suppressive ART (years)** | **HIV DNA copies/10<sup>6</sup> PBMC on ART** | **ART regimen<sup>A</sup>** |
| --- | --- | --- | --- | --- | --- |
| ZA-001 | Female | 10.9 | 8.80 | 86.3 | AZR/3TC/LPV/r |
| ZA-004 | Male | 2.7 | 7.92 | 23.6 | AZR/3TC/LPV/r |
| ZA-006 | Female | 9.0 | 7.45 | 46.7 | AZR/3TC/LPV/r |
| ZA-011 | Female | 9.3 | 7.35 | 181.5 | AZR/3TC/LPV/r |
| ZA-012 | Male | 5.1 | 8.60 | 11.9 | AZR/3TC/LPV/r |

<sup>A</sup> zidovudine (AZR), lamuvidine (3TC), ritonavir-boosted lopinavir (LPV/r)

**Table 2.**
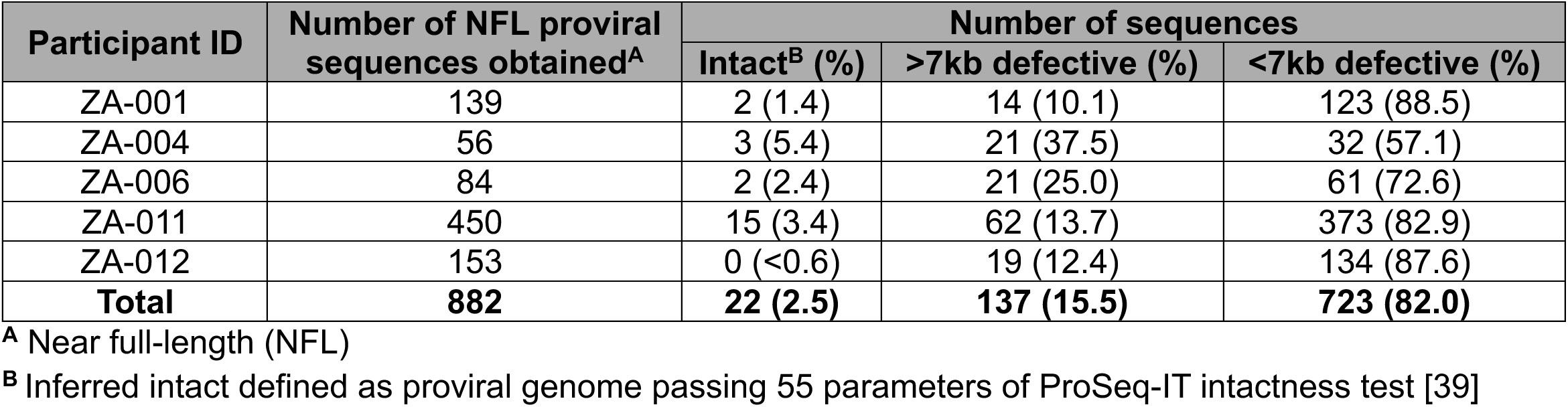
Number of intact and defective proviruses obtained from donors in the CHER cohort.

**Table 3.** Donor characteristics of previously published studies in HIV persistence in infants, children, and adults.

| Study | Age group | Subtype | Donor (n) | Median time of ART initiation (range)<sup>A</sup> | Median time on suppressive ART (range)<sup>A</sup> |
| --- | --- | --- | --- | --- | --- |
| Ho *et al.* [35] | Adult | B | 8 | 73.5m (6m-141m) | 68m (28m-108m) |
| Bruner *et al.* [33] | Adult | B | 10 | 3.28m (17d-6m) | 106.5m (10m-203m) |
| Hiener *et al.* [34] | Adult | B | 6 | 9m (6m-12m) | 125.4m (38.4m-212.4m) |
| Garcia-Broncano *et al.* [38] | Infant<sup>B</sup> | C | 10 | 2.39d (0.20h-4.77d) | 12.5m (1m-24m) |
|  | Children<sup>C</sup> | C | 10 | 7.15m (2.6m-11.7m) | 23.5m (16m-31m) |

<sup>A</sup> Hours (h), Days (d), Months (m)
<sup>B</sup> Early Infant Treatment (EIT) Study: EIT pre-ART and EIT on ART
<sup>C</sup> EIT control group, Delayed ART (EIT Delayed-ART)

We found that children on long-term ART have significantly fewer sequence-intact proviruses than children on short-term ART who received either early or delayed treatment (2.5% vs. 19.6% and 6.1%, p<0.0001) and have fewer intact proviruses than adults on ART analyzed with a similar method (2.5% vs. 5.9%) (p=0.0002) (**Table 4**). Of the defective proviruses in the CHER children (long-term ART), 723 (82%) contained large internal deletions (proviruses <7kb), slightly higher than reported for adults on ART (71%) (p=0.046), and greater than the fraction with large internal deletions in children on short-term ART (46-61%) (p<0.0001) or pre-ART (19%) (p<0.0001) (**Figure 1**). Of the pre-ART proviruses obtained from infants in the EIT study, the majority (55%) were inferred intact. Defective proviruses resulting from mutations or small deletions comprised 16% of sequences in the CHER cohort (children on long-term ART) compared to 34% in children on short-term ART (p<0.0001) and 23% in adults on ART (p=0.0012). This finding suggests a strong selection with time against proviruses in children that are >7kb in length, regardless of whether they are intact or defective.

**Figure 1.**
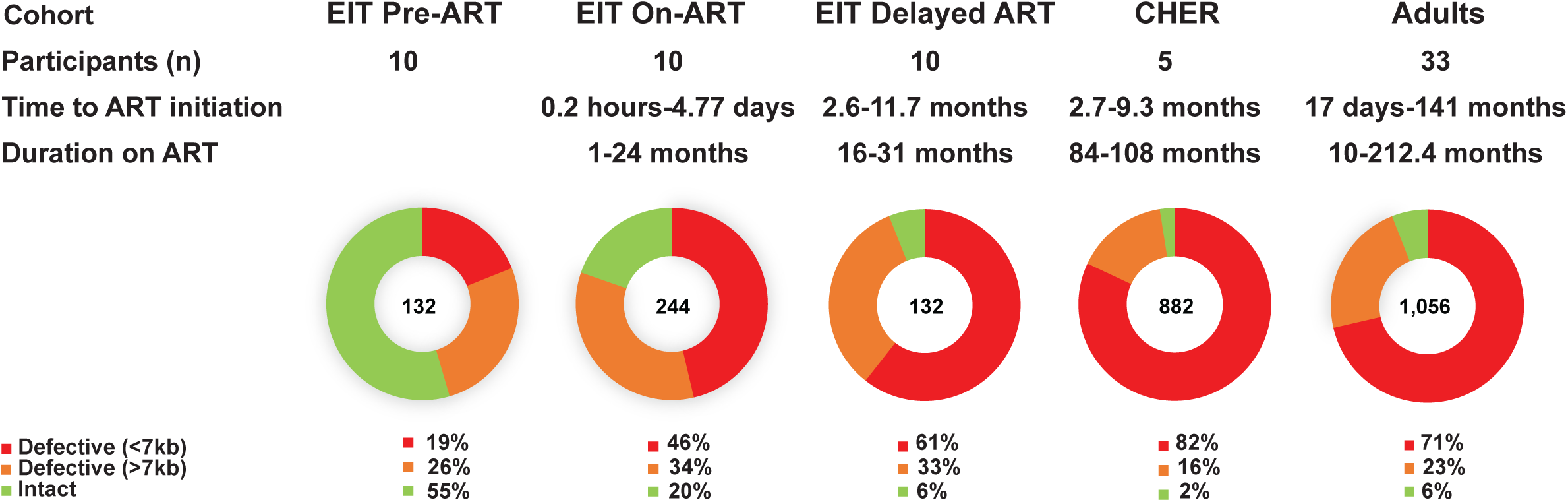
The fraction of intact and defective proviruses in children and adults. Red-Defective proviruses with large deletions (<7kb). Orange-Defective proviruses with lethal mutations and/or small deletions (>7kb). Green-Intact proviruses.

**Table 4.** Number of intact and defective proviruses obtained from donors across cohorts.

| Group<sup>A</sup> | Number of NFL proviral sequences obtained<sup>B</sup> | Number of sequences |  |  |
| --- | --- | --- | --- | --- |
|  |  | Intact<sup>C</sup> (%) | >7kb defective (%) | <7kb defective (%) |
| EIT Pre-ART | 132 | 72 (54.5) | 35 (18.0) | 25 (26.5) |
| EIT On-ART | 244 | 48 (19.7) | 83 (34.0) | 113 (46.3) |
| EIT Delayed-ART | 132 | 8 (6.1) | 44 (33.3) | 80 (60.6) |
| Adults | 1,056 | 63 (6.0) | 239 (22.6) | 754 (71.4) |
| CHER | 882 | 22 (2.5) | 137 (15.5) | 723 (82.0) |

<sup>A</sup> EIT Pre-ART, EIT On-ART, EIT Delayed-ART [38], Adults [33-35], CHER (this study)
<sup>B</sup> Near full-length (NFL)
<sup>C</sup> Inferred intact defined as proviral genome passing 55 parameters of ProSeq-IT intactness test [39]

### Partial or complete env deletions were found in most defective proviruses regardless of ART status or duration

Further investigation of the defective sequences with large internal deletions (proviruses <7kb) revealed that the majority contained complete or partial deletions of the *env* gene in all datasets, regardless of ART status or duration (**Figure 2**). Proviruses with completely deleted *env* genes comprised 40-56% of all sequences <7kb. Large internal deletions that spanned both the 5′ and 3′ halves of the genome, including partial/complete deletions of *env*, comprised 32-52% of proviruses <7kb, while deletions in only the 5′ half of the genome were seen in 3-12% of proviruses <7kb (p<0.0001). These data demonstrate that there is a strong selection pressure against the maintenance of the *env* gene and that this selection pressure exists prior to ART and during short-term and long-term ART. Of the proviruses with large internal deletions that persist on ART, about 90% (blue + yellow; **Figure 2**) include at least a partial deletion of the *env* gene, suggesting *env* confers a deleterious effect on infected cell clone survival, independent of the intactness of the proviral genome.

**Figure 2.**
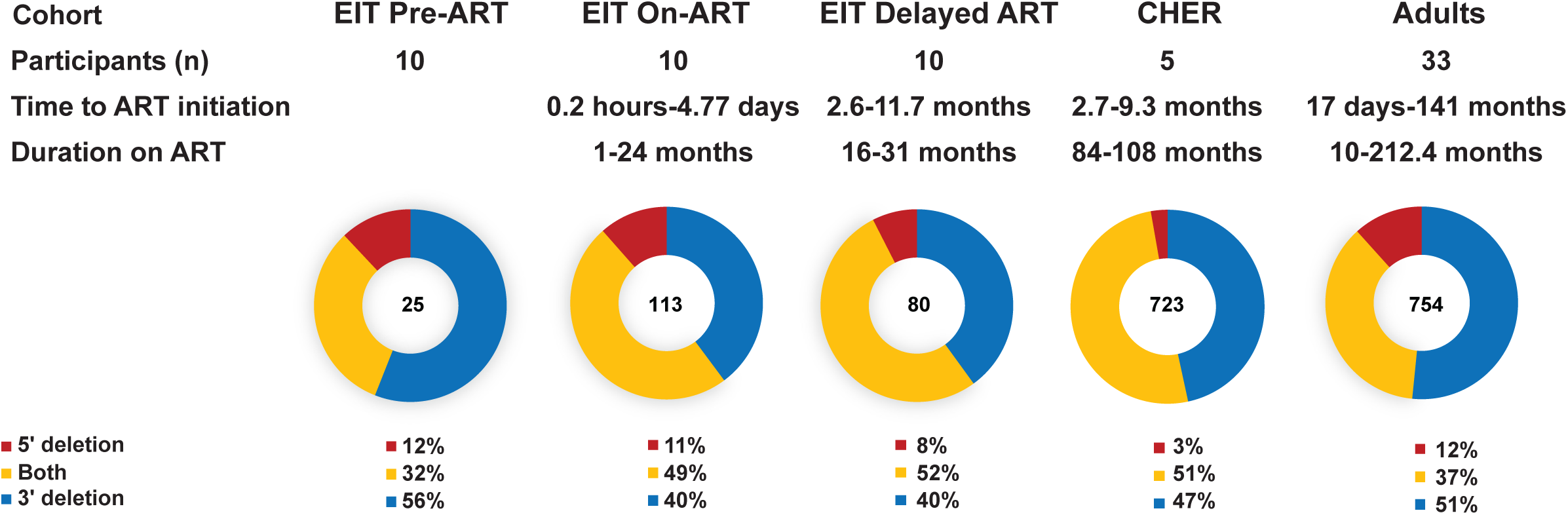
Proviruses with large internal deletions typically include deletion of the *env* gene. Blue- Proviruses with 3′ large internal deletions. Yellow- Proviruses with large internal deletions spanning 5′ and 3′ ends. Red- Proviruses 5′ large internal deletions.

### Most proviruses >7kb in children on long-term ART contain multiple fatal defects

All proviruses that were determined to be defective using the ProSeq-IT tool but did not contain large internal deletions (n=137 sequences), were analyzed for their number defects (**Figure 3**). While most of the proviruses in this subcategory collected prior to ART contained only one defect (51%), most proviruses in the long-term, on-ART datasets contained two or more defects (>81%) (EIT pre-ART vs. CHER p<0.0001 or vs. Adults p<0.0001). The highest variance in observed defects was seen in the EIT on-ART dataset where some proviruses had up to 9 defects. These data show that proviruses with single defects are selected against with time on ART, leaving persistent proviruses with multiple defects in both children and adults on long-term treatment.

**Figure 3.**
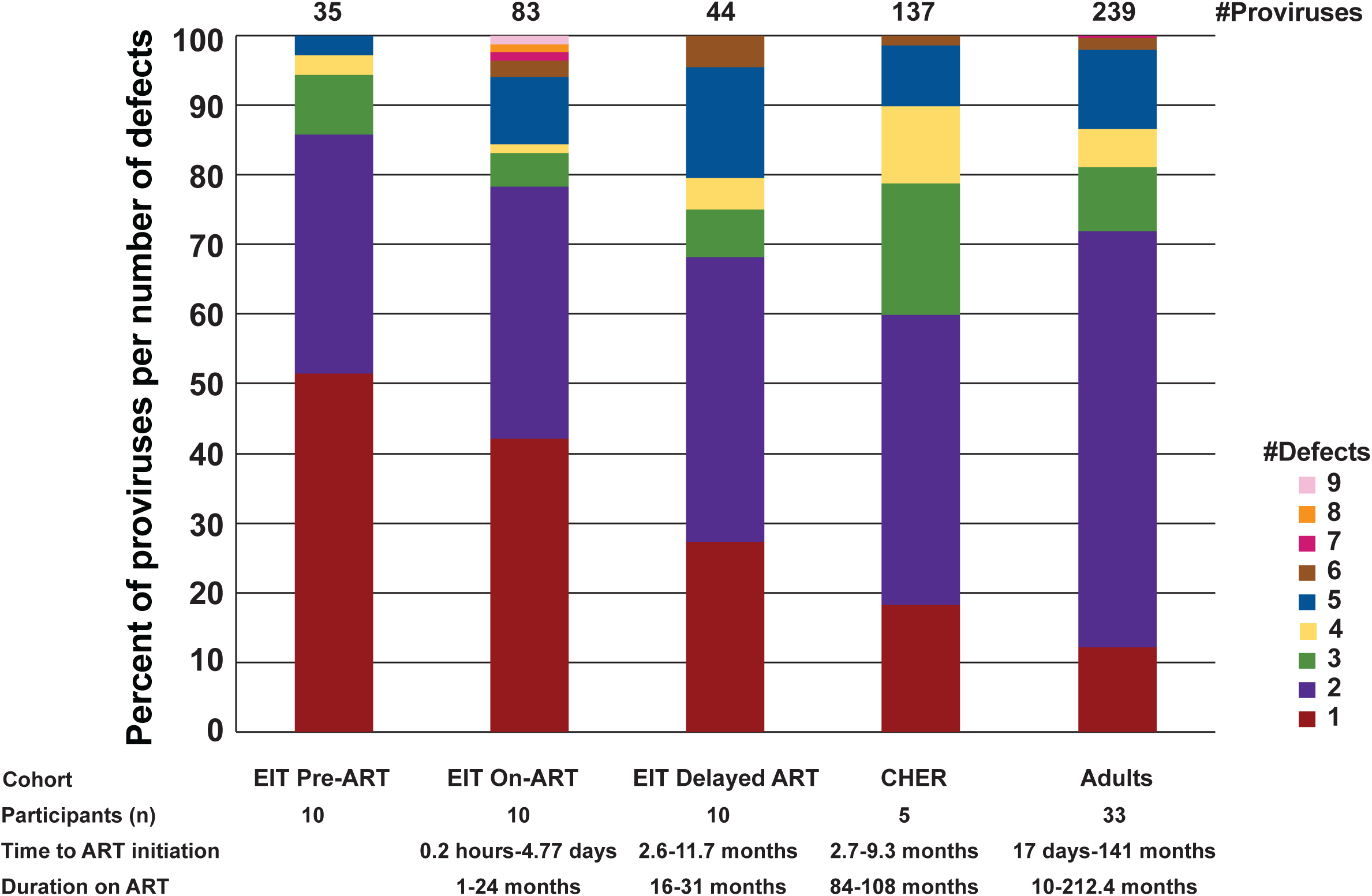
Number of defects in proviruses >7kb. All proviruses that did not have large internal deletions but were not sequence intact were analyzed for the number of different defects. The different colors indicate the number of proviruses with 1, 2, or more defects. The defects analyzed include MSD defect, packaging signal defects, *gag* start codon defect, insertions (*gag*-*pol*), small deletions (*gag*-*pol*), insertion (*env*), small deletion (*env*), frameshift premature stop codons, non-frameshift premature stop codons, and RRE defect.

### Different defects in proviruses that are >7kb in children vs. adults on ART

Of the 137 defective proviruses that were >7kb in the CHER cohort, frameshift-induced stop codons, stop codons in the absence of frameshifts, and *gag* start codon mutations were the most common defects, consistent with the defective proviruses in all infant and children cohorts (**Figure 4**). However, while mutations in the major splice donor site (MSD, red) and packaging signal (Psi, orange) defects were very common in adults on ART (∼38%), they were less frequent in children before and on both short-and long-term ART (<21%) (p<0.0001). The lower frequency of Psi defects in children on ART may influence results of the Intact Proviral DNA assay (IPDA) [40] by having a higher percentage of proviruses that are Psi+. The IPDA is a high throughput method used to estimate the number of intact proviruses in a sample using the presence of Psi and Rev response element (RRE) sequences as a surrogate for NFL sequencing, since Psi and RRE are commonly deleted in proviruses that persist on ART. Although a higher fraction of defective proviruses >7kb in children on ART retain the packaging signal compared to adults on ART, most proviruses contain large internal deletions, including deletions of RRE, preventing these proviruses from scoring positive by IPDA. Another observed difference between persistent proviruses in children on ART compared to adults on ART, is the higher frequency of insertions in children that render the proviruses defective (2.5% vs. 0.7%) (p=0.021). In total, we observed the accumulation and persistence of proviruses with multiple defects with time on ART and we observed varying frequencies regarding types of defects in children compared to adults.

**Figure 4.**
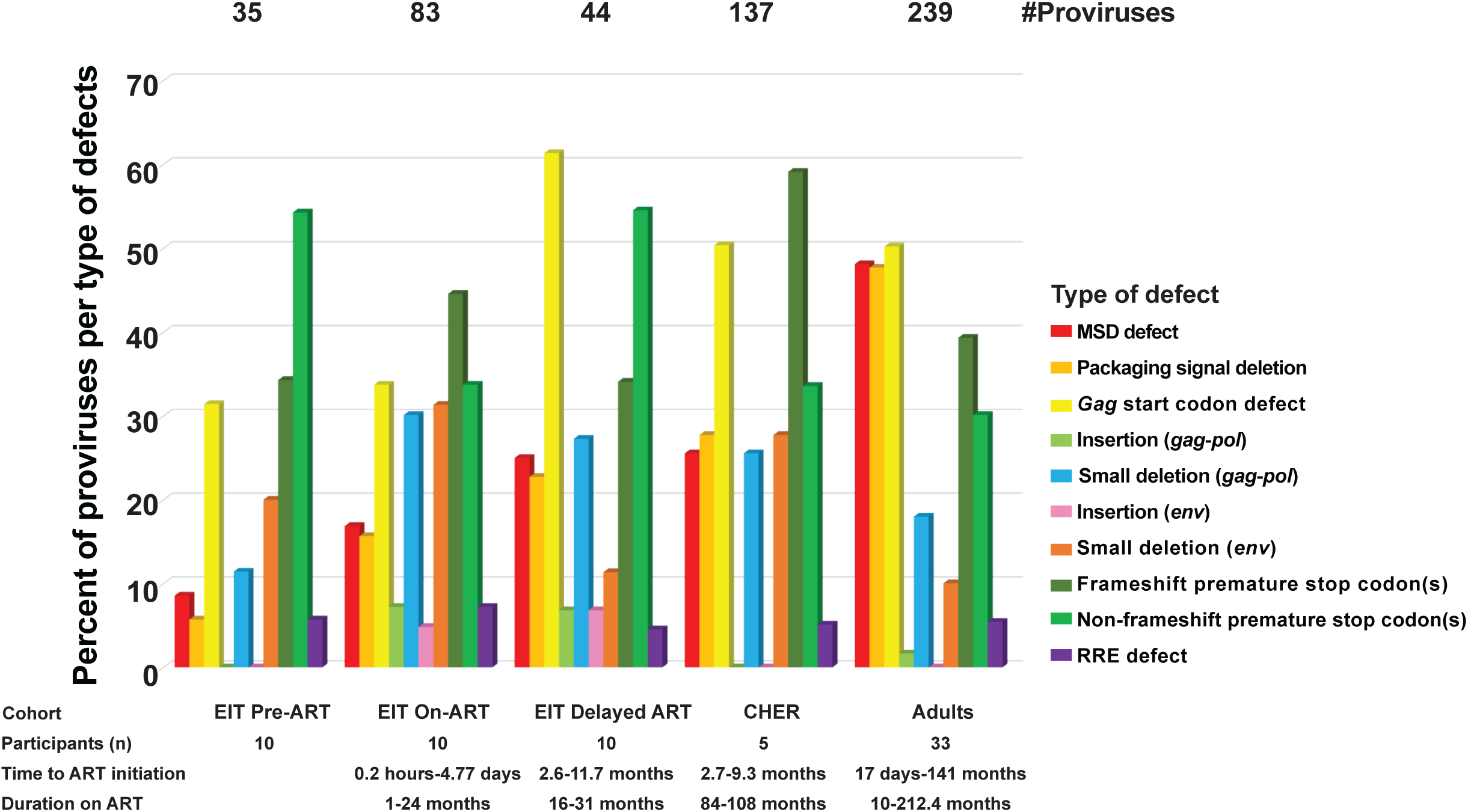
Different types of defects in proviruses >7kb. Red-MSD defect. Orange- Packaging signal defect. Yellow- *gag* start codon defect. Light green- Insertions (*gag*-*pol*). Blue- Small deletions (*gag*-*pol*). Pink- Insertion (*env*) Dark Orange- Small deletion (*env*). Dark green- Frameshift premature stop codons. Green- Non-frameshift premature stop codons. Purple- RRE defect.

### Defects of tat are observed in proviruses that retain env in children

*Tat/rev* coding regions have overlap with the *env* gene in the 3′ region of the HIV genome. We analyzed defective proviruses that retained *env* for possible *tat* and *rev* defects (**Figure 5**). In the EIT pre-ART cohort, most “*env*-retaining” proviruses contained inferred-intact *tat* genes (71%), while in the EIT on-ART cohort, defects in *tat* were observed in 65% of the *env*-retaining proviruses (p<0.0001) (**Figure 5A**). Interestingly, the loss of *tat* did not correlate with time on ART, with children on long-term ART (CHER) having only 40% of *env*-retaining sequences being defective in *tat*. This difference may reflect the large number of proviruses that delete the *env* gene entirely, and therefore, render *tat* defective as well. Surprisingly, we found no correlation between ART treatment and defective *rev* genomic regions in the *env*-containing proviruses. Instead, we found that *rev* was retained in 50% of env-containing proviruses in children both before and during ART (**Figure 5B**). Retention of *rev* in on-ART cohorts may allow for vRNA/protein production, which should be a target of future studies towards achieving HIV-1 remission without ART. Unique visibility of such epitopes, even on ART, may present an opportunity for immunotherapy to boost reservoir elimination.

**Figure 5.**
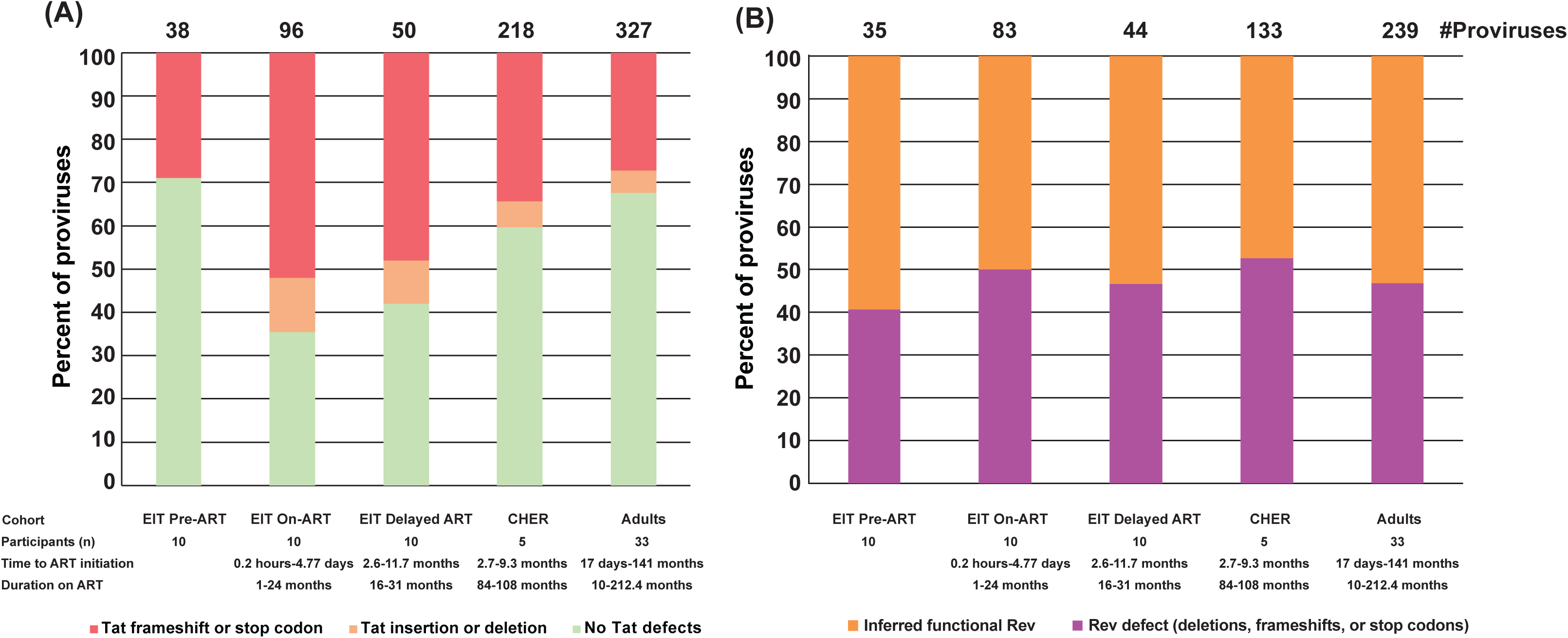
*Tat and rev* characterization of env-retaining defective proviruses. (**A**) *Tat* characterization of env-retaining defective proviruses (**B**) *Rev* characterization of env-retaining defective proviruses

## DISCUSSION

Understanding the impact of the timing of ART initiation and the duration of suppressive ART on persistent provirus genetics is important for defining targets for curative strategies, especially in children with perinatally acquired HIV-1 who face a lifetime on ART. Previous studies have characterized the proviral reservoirs of infants and children, however, the defective proviral landscape on long-term ART was not fully profiled [32-35, 38]. In this study, we performed a deep characterization of the proviral landscape in children in the CHER cohort (treated within 2.7-10.9 months of age) and on long-term ART (84-108 months), compared to previously published datasets of children with differing times of ART initiation and duration: early treated infants (EIT) (treated within 0.2 hours - 4.77 days) on short-term ART (1-24 months) and infants who initiated ART at similar ages to those in the CHER cohort (treated within 2.6-11.7 months) but on short-term ART (16-31 months) [38]. We also compared proviral landscapes in the children in the CHER cohort to those in adults on ART (10-212 months) [33-35].

In the samples from the CHER cohort (long-term ART), the majority of proviruses were defective, and only 2.5% were found to be intact. This finding is consistent with previous data from children [32], but is lower than the fraction of intact proviruses reported for adults on ART using a similar approach (6%) [33-35]. In contrast, in the “EIT pre-ART” dataset, most proviruses were inferred intact, as is expected in uncontrolled infection. In the on-ART datasets, many-to-most defective proviruses contained large internal deletions regardless of the time of ART initiation or the duration of ART. Of the on-ART datasets, the “EIT on-ART” group (short-term ART) had the smallest fraction of sequences with large internal deletions, consistent with the shortest duration on ART. This fraction was higher in the CHER children with longer durations on ART, suggesting a selection against proviruses that retain internal genes, as previously seen in adults [41, 42]. In all datasets, large internal deletions with partial or complete deletion of 3′ genes were most commonly observed, potentially resulting from template switching during reverse transcription. *Env*-deleted genomes have also been commonly observed in studies investigating the proviral landscapes in adults [34, 35, 43, 44], suggesting that *env* expression is highly selected against in the persistence of infected cells on ART in both children and adults [45]. A recent study by Botha *et al.* found that the largest infected cell clones in children on long-term ART contain only solo LTRs [37], suggesting that the decline of intact proviruses may be due, in large part, to homologous recombination of the LTRs post-integration. This observation aligns with the mechanism of solo LTR formation of human endogenous retroviruses (HERVs), as solo LTRs comprise as much as 90% of the HERVs arising from homologous recombination of endogenous LTRs [46-49]. Although the NFL single-genome sequencing approach used here does not capture solo LTR proviruses, combined with the previous study by Botha *et al.* which was performed on the same cohort of children, our results suggest that >90% of proviruses that persist in children for 7-9 years on ART contain large internal deletions of either the *env* gene or of all internal HIV-1 genes.

In the proviruses retaining the *env* gene, the *rev* coding region was inferred to be functionally intact in about 50% in both children and adults, likely due to the flexibility of several structural features of Rev [50-54]. This finding is consistent with a study from Imamichi *et al.* which reported that some defective proviruses can produce viral proteins that elicit CTL responses, suggesting that HIV unspliced and partially spliced products from defective proviruses are exported from the nucleus in a Rev-dependent manner [50]. Unlike *rev*, *tat* defects appeared to be selected during ART and potentially more so in children than in adults [55]. However, it is important to consider that >85% of all proviruses that persist on ART have large internal deletions that render both *tat* and *rev* defective (not including the solo LTRs). As *tat* and *rev* function in viral expression and *env* encodes a surface protein, these data strongly suggest that the ability to express genes in the 3’ region influences the selection of proviruses by immune pressures, such as CTL and/or neutralizing antibodies [45, 56-63]. The fact that the splice acceptor sites required for both the completely spliced *tat*/*rev* and the partially spliced *vpu*/*env* viral RNA species are located in overlapping regions (A4cab/A5 and A7) and, since the sequences of the *vpr*-*tat*/*rev* exon 1 and the *env*-*tat*/*rev* exon 2 containing major splicing elements (such as donor and acceptor sites, enhancers, and suppressors for introns/exons) are relatively close to the *env* gene, a provirus retaining *env* might also maintain *tat*/*rev* expression and downstream splicing. Consequently, these gene products may be good targets for engineered antibodies or T cells designed to achieve HIV-1 remission without ART [64, 65], possibility in combination with latency reversing agents.

The “EIT delayed-ART” group, the CHER children, and the adults all had high frequencies of defects in the *gag* start codon, suggesting that the failure to express *gag* proteins may be favorable for persistence of infected cells on ART. However, the children had lower frequencies of Psi deletions compared to adults. It is not clear if this difference is related to the differential immune systems of children vs. adults or to the different HIV-1 subtypes in the cohorts studied (HIV-1 subtype C in the children vs. subtype B in the adults). IPDA, and similar assays, are useful tools to differentiate intact proviruses from defective proviruses using primers and probes that target the Psi and RRE regions of the HIV-1 provirus. A positive IPDA signal indicates an intact provirus [40, 66]. Although larger studies are needed, the more frequent retention of Psi in defective proviruses in children on ART suggests that this target may be a less appropriate for measuring intact proviruses in children with subtype C infection. Measuring the HIV-1 reservoir in children with subtype C infection may more appropriately target defects in the *gag* start codon which appear to be more frequent than defects in Psi. Although the RRE was not frequently defective in the proviruses not containing 3’ deletions, it is important to note that >85% of the total proviruses in children on ART contained 3’ deletions, including RRE, making RRE an appropriate IPDA target for children, as it is in adults. Based on these results, alternative screening methods in which probes targeting multiple targets, such as Q4PCR [67] or Rainbow proviral HIV-1 DNA dPCR [68], may be more informative.

A comparison of pipelines used in proviral sequence analysis reported bias in the characterization of defective sequences [69]. As there is no consensus pipeline to classify defective sequences, variance in reported defects leads to an imprecise record of the proviral landscape of people on ART. Furthermore, some pipelines use an order of elimination, in which sequences are binned into defective categories and excluded from further investigation, leading to an underestimation of the frequencies of some defects and an overestimation of others [69]. Here, we used the ProSeq-IT tool which identifies all known defects in proviral sequence datasets [39]. Using this approach, we found the accumulation of multiple defects in proviruses that persist in both children and adults on ART, with some proviruses containing up to 9 different lethal mutations. This observation may be due to limited expression of proviruses with multiple mutations while some single mutations may be tolerated and permit expression and detection of viral RNA and proteins. The accumulation of multiple defects may reflect selection against transcriptional/protein production of any HIV genes, thus leaving highly defective, transcriptionally mute proviruses.

Although more than 50% of the proviruses in the pre-ART dataset were inferred intact, stop codons, likely from APOBEC3G-derived hypermutation, was the most frequent defect, suggesting that APOBEC3G may contribute to pre-ART control of replication in children. On-ART, the children treated in infancy contained the highest frequency of proviruses with multiple fatal defects as well as the most variance in defect type. This finding suggests that the proviral landscape undergoes the most change early after ART initiation as infants experience a rapid decline of HIV-1 DNA due to culling of infected cells expressing viral proteins. This decline of infected cells results in a shift of the proviral population to those with a wide variety of defects [70]; consistent with the Koofhethile *et al.* observation of rapid decline in intact proviruses within one month of treatment initiation [27]. These findings suggest that the time of ART initiation may play a significant role in shaping the proviral landscape. Guidelines are in place to rapidly treat infants born to mothers with HIV-1; however, adults living with HIV-1 may experience longer durations between initial infection and ART initiation. This difference in the duration of uncontrolled infection prior to ART initiation may play a key role in the differences we observed between the proviral landscapes of children vs. adults, in addition to the fact that the mature immune system of adults may differentially influence the evolution of the reservoir compared to the naivety of the infant immune system.

While this study adds to the limited data on children living with HIV, low sample availability and low proviral loads are a major limitation to generating sequence data. More donors/participants are required. The method of amplicon generation also influences the characterization of the proviral landscape, as recent comparisons of amplification methodologies demonstrated a bias for different sized amplicons [71]. Additionally, our NFL amplification approach precludes the detection of solo LTRs. New methods are needed to capture both solo LTRs and proviruses containing internal genetic information to fully characterize proviral landscapes on ART. Nonetheless, we highlight important differences in the children vs. adult proviral landscape, especially in the more frequent retention of intact MSD and packaging signal in children on short and long-term ART, and the apparent stronger selection against *tat* retention in children on short-term ART. These findings highlight prime targets for the future design of high throughput assays to measure HIV reservoir size in children and may present opportunities for new strategies to control HIV infection unique to pediatric cohorts.

## MATERIALS AND METHODS

### Donor samples

PBMC were obtained from 5 children living with HIV-1 and on long-term ART (7.4-8.8 years) who were enrolled in the CHER trial (NCT00102960) (**Table 1**). Further demographics and virologic characteristics of these children have been reported in van Zyl *et al.* [31], Katusiime *et al.* [32], and Bale *et al.* [36]. The children had their viremia fully suppressed on ART for the duration of treatment, had no evidence of ongoing cycles of viral replication and contained large, infected cell clones with solo HIV LTRs. The legal guardians provided informed consent for PBMC collection and for sequencing of HIV proviruses and integration sites. The study was approved by the internal review board at Stellenbosch University.

### Genomic DNA extraction

Genomic DNA (gDNA) was extracted from 1.25 × 10^6^ PBMC from each child. In brief, cell pellets of 1.25 × 10^6^ PBMC were subjected to guanidine HCl (Sigma-Aldrich: Cat#G9284) lysis, cleaned by isopropanol/ethanol precipitation, and resuspended in 5mM Tris-HCl (pH 8.0) as described in Katusiime *et al.* [32]. The resuspended DNA was incubated at 42°C for 2 hours then stored at -80°C prior to undergoing near full-length HIV PCR amplification and sequencing as described below.

### Near full-length HIV single-genome sequencing (NFL-SGS)

Extracted gDNA was diluted to an endpoint (0-1 proviral genomes per well, approximately 10 HIV-positive wells per 96-well plate) with 5 mM Tris-HCl (pH 8.0). Near full-length (NFL) PCR was performed on each well and was followed by nested PCR (**Figure S1**). The forward PCR primers were designed in 5′ LTR_U5 and *gag* leader and the reverse primers were designed in U5 of the 3′ LTR (**Fig. S1, Table S1**). NFL amplification was initially performed using Ranger mix (Bioline: Cat#BIO-25052) for the material from donors ZA-004, ZA-006, and ZA-011. A 200μL volume of diluted gDNA was added to a master mix comprised of 500μL 2x Ranger Mix, 284μL nuclease free water, and 8μL each of 50μM forward and reverse primer. The total volume was spread across a 96-well plate (10μL per well). PCR cycling conditions are described in **Table S2**. NFL amplification from donors ZA-001 and ZA-012 was performed using Platinum SuperFi II DNA polymerase (Thermofisher Scientific: Cat#12361010). A master mix containing 210μL 5x SuperFi buffer, 20μL of 10μM dNTP mix, 20μL each of 10μM forward and reverse primers, 559.5μL nuclease free water, and 10.5μL SuperFi II DNA polymerase was prepared. 220μL of diluted gDNA was spread across a 96-well plate (10μL per well) along with the master mix and was PCR amplified as in **Table S2**. Invitrogen 96 well 1% agarose E-Gels (Thermofisher Scientific: Cat#G700801) were used to identify positive wells. Amplicons were then run on 12 well 0.8% agarose E-Gels (Thermofisher Scientific: Cat#G501808) along with a 1kb plus DNA ladder (Thermofisher Scientific: Cat#10488090) to determine the amplicon size.

### Illumina sequencing

The resulting amplicons were sequenced by next generation sequencing (NGS) using the Illumina MiSeq platform. Due to low fidelity 5′ phosphorylation by the Illumina preparation kit, a single round of PCR was performed using phosphorothioated primers. The amplicons were then purified using AMPure XP beads (Beckman Coulter: Cat#A63881) according to the manufacturer protocol and were quantified via FluoroSkan. Sequencing was done in multiplexed libraries using the NEBNext Ultra II FS DNA Library Preparation Kit and protocol (New England Bio Labs: Cat#E7805L). In brief, the amplicon DNA was fragmented, blunted, and end repaired with 5’ phosphorylation and 3′ dA-tailing. Hairpin loop adapters were ligated to the fragments and USER enzyme was used for U excision to free the adapter ends. The sample was enriched for adaptor ligated fragments of 150-250bp via size exclusion AMPure XP bead purification. The libraries were enriched by PCR and barcoded to allow for multiplexed libraries. The libraries underwent a final PCR cleanup with AMPure XP bead purification and were assessed for quality using TapeStation. Pooled libraries were run on the Illumina MiSeq using 300 cycle MiSeq Reagent Nano Kit v2 (Illumina: Cat#MS-103-1001).

### Sequence analysis

Resulting Illumina sequences were assembled and aligned using the MAFFT alignment program via online server [72]. The Proviral Sequence Annotation & Intactness Test (ProSeq-IT) tool (https://psd.cancer.gov) was used to annotate defects throughout the provirus and to infer genetic intactness [39]. The tool infers intactness based on 19 parameters, including sequence length, major splice donor (MSD) site, packaging signal, *gag* start codon status, rev response element (RRE) region status, insertions, deletions and premature stop codons in *gag*, *pol*, and *env*. The fraction of intact proviruses and those with major internal deletions or mutations in each donor was determined. Proviral populations were also analyzed based on the types and abundance of defects (**Figure 1**). The fraction of proviruses with defects or deletions in *tat* and *rev* were also determined. To infer “functional” Rev, a multiparameter approach similar to ProSeq-IT was used. Some considerations included the 18Q/60R alpha-helix stability core sites, the M10 dominant-negative mutant (L78D/E79R), presence of premature stop codons or frameshifts, and deletions within the first 58 amino acids of exon 2 or a deletion of the first 1-10 amino acids in exon 1 [53, 54, 73]. Due to the multiple subtypes present in the cohorts examined, subtype B/D typically exhibit the 21bp deletion in the C-terminus (‘TQQSPGT’ motif) relative to other HIV-1 subtypes. Further, it is common to observe the transition mutation of C®T (<u>C</u>AA®<u>T</u>AA; Glutamine®Stop) in subtype C in the C-terminus. These analyses were also performed on previously published datasets for comparison to our dataset. The previously published datasets were obtained from the HIV Proviral Sequence Database (PSD) [74], and include infants treated <5 days after birth (Early Infant Treatment Study (EIT)) [38], children on short-term ART (<2.5 years) (EIT delayed-ART) [38], and adults on ART for a median of 9.26 years [33-35]. More details on the previously published datasets are provided in **Table 2**.

### Statistical analysis

All statistical analyses were performed using R (version 4.3.1). If the analysis compared two disjoint sets (*e.g.*, defective versus intact) and only two cohorts, a Fisher exact test was used. If there were more than two cohorts in the comparison, a Chi-square test was used. For a comparison of two groups where a clear increase or decrease was observed, a Binomial test was used. Note that the Fisher and Chi-squared tests were run as 2-tailed tests, while the Binomial test was a 1-tailed test.

## AUTHOR CONTRIBUTIONS

Conceptualization- JMH, MGK, MFC, GVZ, SCP, MFK; Methodology- JMH, MGK, AAC, MJB, BTL, WS; Formal Analysis- JMH, AAC, MJB, BTL.; Original Draft Preparation- JMH.; Review & Editing- JMH, MGK, AAC, MJB, BTL, WS, MFC, GVZ, SCP, and MFK.; Visualization- JMH, SCP, MFK.; Supervision- SCP and MFK.

## FUNDING

This research was supported in part by the Intramural Research Program of the NIH. This work was supported by intramural NCI funding (ZIA BC 011697) to the HIV Dynamics and Replication Program (MFK) and by the Office of AIDS Research. This study was supported by funding from the National Institutes of Allergy and Infectious Diseases (Comprehensive International Program for Research on AIDS [CIPRA]–South Africa, grant U19 AI153217); the U.S.-South Africa Program for Collaborative Biomedical Research; the National Cancer Institute (grant U01CA200441). Other funders include NCI contract No. 75N91019D00024 to BTL, Leidos Biomedical Research, Inc. The content is solely the responsibility of the authors and does not necessarily represent the official views of the National Cancer Institute, National Institutes of Health, or the Department of Health and Human Services.

## DATA AVAILABILITY

All NFL sequences have been deposited in NCBI (PV625462-PV626342) and the NCI Proviral Sequence Database (*in process*) using the PubMed ID of this manuscript. Additional data availability requests can be fulfilled by contacting the corresponding author(s).

## ACKNOWLEDGEMENTS

We thank the participants and their caregivers for their willingness to take part in this study, as well as the clinic staff for their assistance. We acknowledge our collaborative interactions with the Behavior of HIV in Viral Environments Center (U54AI170855). We thank the entire HIV Dynamics and Replication Program administrative staff.

## CONFLICTS OF INTEREST

The authors have no conflicts of interest.

## SUPPLEMENTAL TABLES AND FIGURES

**Table S1.**
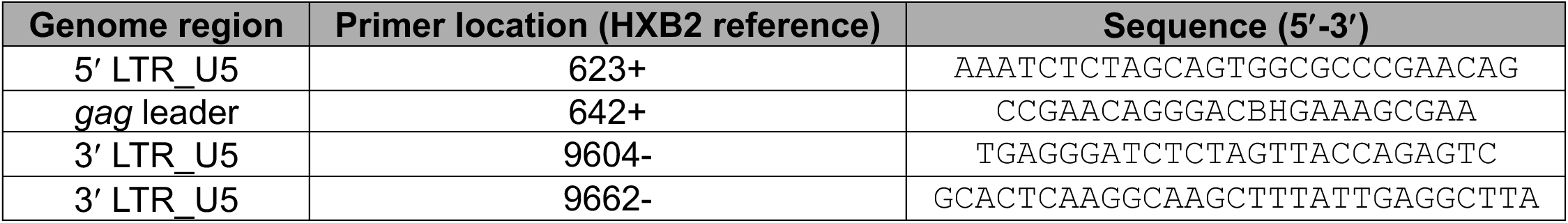
PCR primers used for near full-length HIV PCR amplification.

**Table S2.** PCR conditions for near full-length sequencing of HIV proviruses.

| **Polymerase** | **PCR1 conditions** | **PCR2 conditions** |
| --- | --- | --- |
| Ranger | 1. 95°C for 2 min<br>2. 98°C for 10 sec<br>3. 61.5°C for 10 min<br>4. 72°C for 10 min<br>5. Go to Step 2, 30x cycles<br>6. 4°C hold | 1. 95°C for 2 min<br>2. 98°C for 10 sec<br>3. 65.5°C for 10 min<br>4. 72°C for 10 min<br>5. Go to Step 2, 30x cycles<br>6. 4°C hold |
| SuperFi II | 1. 98°C for 3 min<br>2. 98°C for 10 sec<br>3. 61.5°C for 30 sec<br>4. 72°C for 5 min<br>5. Go to Step 2, 30x cycles<br>6. 72°C for 5 min<br>7. 4°C hold | 1. 98°C for 3 min<br>2. 98°C for 10 sec<br>3. 65.5°C for 30 sec<br>4. 72°C for 5 min<br>5. Go to Step 2, 30x cycles<br>6. 72°C for 5 min<br>7. 4°C hold |

**Figure S1.**
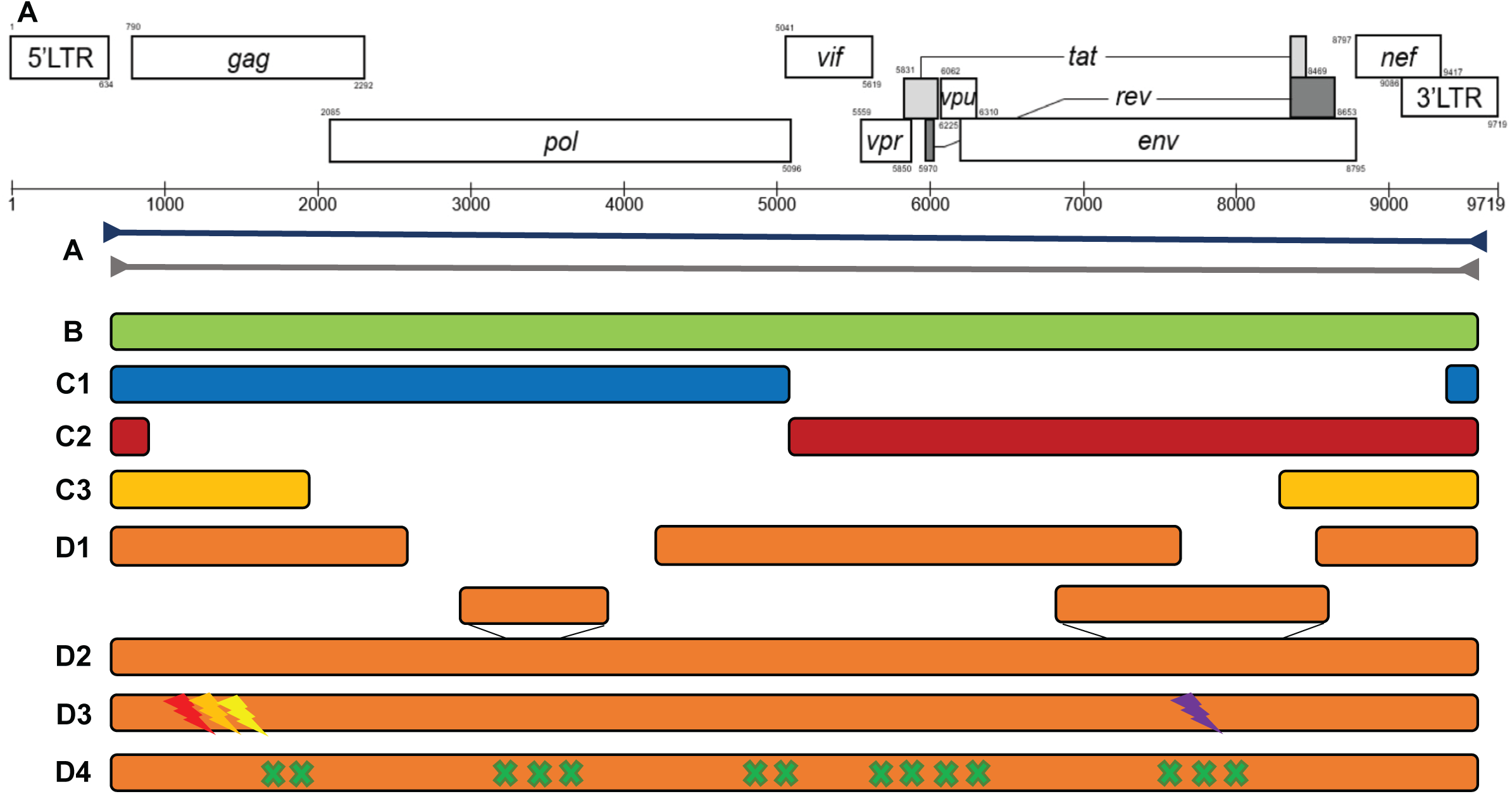
Various proviral structures resulting from HIV-1 near full-length (NFL) amplification (∼9kb). A. Reference HIV-1 genome (accessed from https://www.hiv.lanl.gov). B. Intact HIV-1 genome. C. Genome with large internal deletion. C1. Genome with 3’ deletion. C2. Genome with 5’ deletion. C3. Genome with both 5’ and 3’ deletion. D1. Small internal deletions. D2. Insertions. D3. Defects in regulatory elements including major splice donor site, packaging signal, *gag* start codon, *rev* response element. D4. Premature stop codons resulting from hypermutation.

## Notes

### Competing Interest Statement

The authors have declared no competing interest.

